# Decreased adaptation at human disease genes as a possible consequence of interference between advantageous and deleterious variants

**DOI:** 10.1101/2021.03.31.437959

**Authors:** Chenlu Di, Diego Salazar Tortosa, M. Elise Lauterbur, David Enard

**Author notes:** **Corresponding author** David Enard.

## Abstract

Advances in genome sequencing have dramatically improved our understanding of the genetic basis of human diseases, and thousands of human genes have been associated with different diseases. Despite our expanding knowledge of gene-disease associations, and despite the medical importance of disease genes, their evolution has not been thoroughly studied across diverse human populations. In particular, recent genomic adaptation at disease genes has not been well characterized, even though multiple evolutionary processes are expected to connect disease and adaptation at the gene level. Understanding the relationship between disease and adaptation at the gene level in the human genome is severely hampered by the fact that we don’t even know whether disease genes have experienced more, less, or as much adaptation as non-disease genes during recent human evolution. Here, we compare the rate of strong recent adaptation in the form of selective sweeps between disease genes and non-disease genes across 26 distinct human populations from the 1,000 Genomes Project. We find that disease genes have experienced far less selective sweeps compared to non-disease genes during recent human evolution. This sweep deficit at disease genes is particularly visible in Africa, and less visible in East Asia or Europe, likely due to more intense genetic drift in the latter populations creating more spurious selective sweeps signals. Investigating further the possible causes of the sweep deficit at disease genes, we find that this deficit is very strong at disease genes with both low recombination rates and with high numbers of associated disease variants, but is inexistant at disease genes with higher recombination rates or lower numbers of associated disease variants. Because recessive deleterious variants have the ability to interfere with adaptive ones, these observations strongly suggest that adaptation has been slowed down by the presence of interfering recessive deleterious variants at disease genes. These results clarify the evolutionary relationship between disease genes and recent genomic adaptation, and suggest that disease genes suffer not only from a higher load of segregating deleterious mutations, but also an inability to adapt as much, and/or as fast as the rest of the genome.

## Introduction

Advances in genome sequencing have dramatically improved our understanding of the genetic basis of human diseases, and thousands of human genes have been associated with different diseases (Amberger et al., 2019; Piñero et al., 2020). Despite our expanding knowledge of gene-disease associations, and despite the fact that multiple evolutionary processes might connect disease and genomic adaptation at the gene level, these connections are yet to be studied. Different evolutionary processes have the potential to make the occurrence of disease genes and adaptation not independent from each other in the human genome. For instance, hitchhiking of deleterious mutations linked to advantageous mutations might increase the risk of disease-causing variants at genes subjected to past directional adaptation. Disease genes might then appear to have experienced more adaptation than non-disease genes if this specific process was sufficiently widespread. Conversely, higher evolutionary constraint, and higher pleiotropy might reduce adaptation at disease genes compared to genes not involved in diseases (Otto, 2004). There is currently considerable uncertainty about how any of these non-exclusive evolutionary processes, or other processes, might have influenced adaptation at disease genes. It is even not well-known whether human non-infectious disease genes have similar, higher or lower levels of adaptation in human populations compared to genes not involved in diseases. Comparing levels of adaptation between disease genes and non-disease genes is a first important step toward better understanding the evolutionary relationship between non-infectious diseases and genomic adaptation.

Multiple recent studies comparing evolutionary patterns between human disease and non-disease genes have found that disease genes are more constrained and evolve more slowly (lower ratio of nonsynonymous to synonymous substitution rate, dN/dS, in disease genes) (Blekhman et al., 2008; Park et al., 2012; Spataro et al., 2017), An older comparison by Smith and Eyre-Waler (2003) found that disease genes evolve faster than non-disease genes (higher dN/dS), but we note that the sample of disease genes used at the time was very limited.

The significant increase of the number of known disease genes since these studies were completed makes it important to update the comparison of evolutionary patterns at disease and non-disease genes. More critically however, past studies all have in common an important limitation that justifies comparing disease genes and non-disease genes again. Disease and non-disease genes may differ by more than just the fact that they have been associated with disease or not. Disease and non-disease genes may also differ in many other factors other than their disease status. Such factors can be a problem when comparing adaptation in disease genes and non-disease genes, because they, instead of the disease status itself, could explain differences in adaptation. For example, disease genes tend to be more highly expressed than non-disease genes (Spataro et al., 2017) (Figure 1). If higher expression happens to be associated with more adaptation in general, one might detect more adaptation in disease genes in a way that has nothing to do with disease, and just reflects their higher levels of expression. Many other factors may also be important. For example, immune genes, which often adapt in response to infectious pathogens, may further complicate comparisons if they are represented in unequal proportions between non-infectious disease and non-disease genes. Comparing genomic adaptation in disease and non-disease genes thus requires careful consideration of confounding factors.

**Figure 1.**
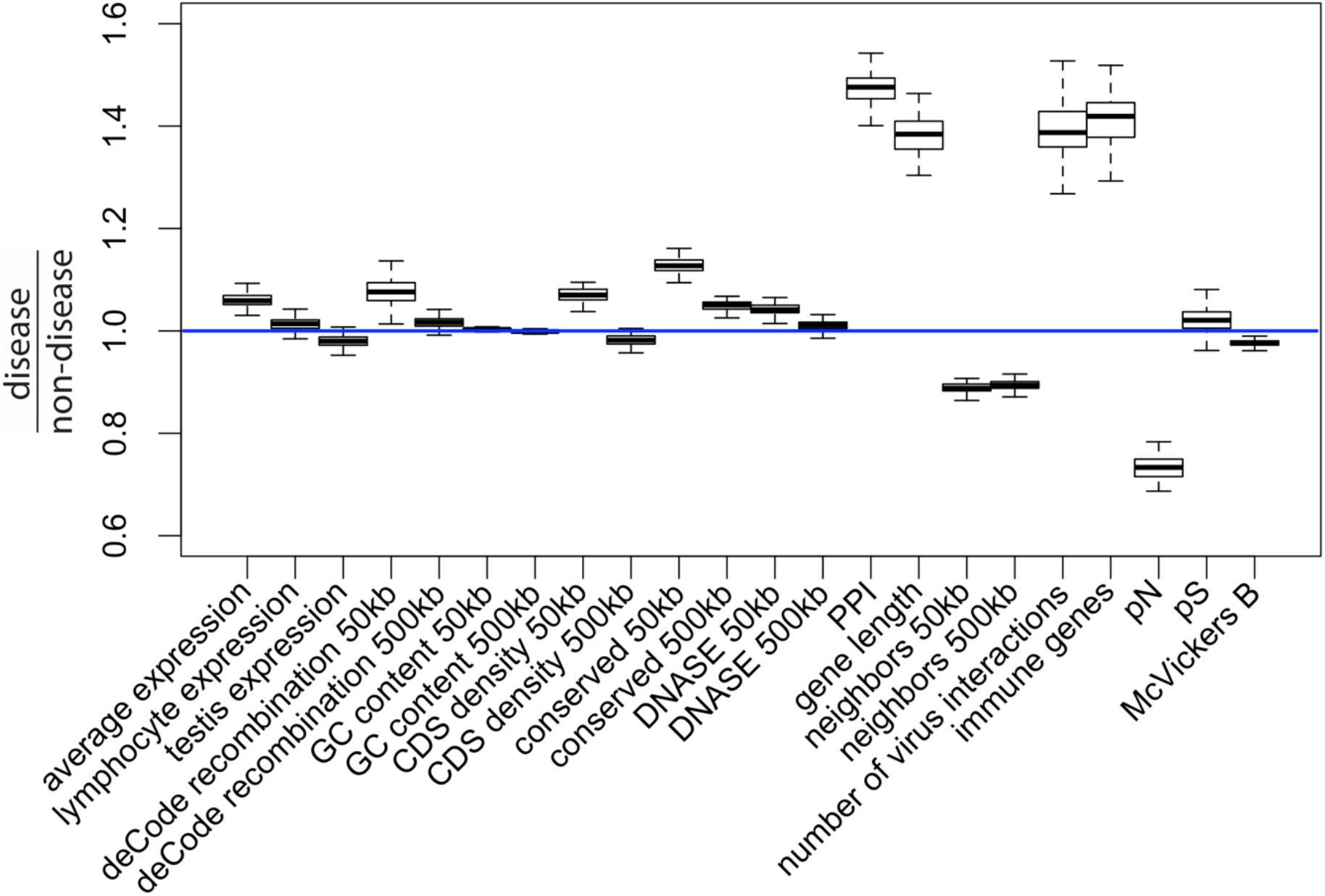
Potential confounding factors in disease versus non-disease genes. Each potential confounding factor is detailed in the Methods. For each confounding factor, the boxplot shows on the y-axis the ratio of the average factor value for disease genes, divided by the average factor value for non-disease genes. The boxplot error bars are obtained by calculating the ratio 1,000 times, each time by randomly sampling as many non-disease genes as there are disease genes.

Among other confounding factors, it is particularly important to take into account evolutionary constraint, i.e the level of purifying selection experienced by different genes. A common intuition is that disease genes may exhibit less adaptation because they are more constrained (Blekhman et al., 2008), leaving less mutational space for adaptation to happen in the first place. Less adaptation at disease genes might thus represent a trivial consequence of varying constraint between genes (Kim et al., 2007), which says little about a specific connection between disease and adaptation. In the same vein, one might expect disease genes to be associated with higher mutation rates, and more frequent adaptation to follow as a trivial consequence of elevated mutation rates. Whether disease genes experience higher mutation rates is however still an open question (Osada et al., 2009; Eyre-Walker and Eyre-Walker, 2014). In any case, focusing specifically on disease and adaptation requires controlling for confounders such as constraint and mutation rate (see Methods, Results and Figure 1 for a complete list of confounders accounted for in this analysis).

A specific evolutionary relationship may exist between adaptation and disease beyond the simple effect of constraint, mutation rate or other confounders. In an evolutionary context, once constraint and other confounding factors have been accounted for, we can imagine three potential scenarios for the comparison of adaptation between disease and non-disease genes. Under scenario 1, any potential difference in adaptation between disease and non-disease genes is entirely due to differences in constraint and other confounding factors. Under this scenario, there is no further evolutionary process linking disease and adaptation together. Therefore, there is no difference in adaptation between disease and non-disease genes once confounding factors have been accounted for.

Under scenario 2, disease genes have more adaptation than non-disease genes. For example, as already mentioned above, deleterious mutations can hitchhike together with adaptive mutations to high frequencies in human populations (Birky and Walsh, 1988; Barreiro and Quintana-Murci, 2010; Chun and Fay, 2011). Other, less well established, cases can be imagined where past adaptation decreased the robustness of a specific gene, and subsequent mutations become more likely to be associated with diseases (Xu and Zhang, 2014). Scenario 2 thus favors a relationship between adaptation and disease, where past adaptation precedes and influences the likelihood of a gene being associated with disease.

Under scenario 3, disease genes have less adaptation than non-disease genes even after accounting for confounding factors such as evolutionary constraint. Such a scenario might occur for example if disease genes happen to be genes that can be sensitive to changes in the environment, with a fitness optimum that can change over time, but where adaptation has not occurred yet to catch up with the new optimum. Such an adaptation lag (or lag load, to reuse the terminology introduced by J. Maynard-Smith (1976)) may occur for example if higher pleiotropy at disease genes (Ittisoponpisan et al., 2017) makes it less likely for new mutations to be advantageous (Otto, 2004) (in addition to increasing the level of constraint already accounted for as a confounding factor). Such an adaptation lag, with genes further away from their optimum, might make such genes more prone to accumulate disease variants that fall too far from the “normal” functioning range around the optimum. An adaptation lag may also occur if deleterious mutations interfere with and slow down adaptation at disease genes more than at non-disease genes (Assaf et al., 2015; Hill and Robertson, 1966).

Even though uncovering the underlying evolutionary processes that govern the relationship between disease and adaptation will take a lot more work than the present analysis, it is important to find first which scenario is the most likely to be true, i.e whether disease genes have as much, more, or less adaptation than non-disease genes. Finding out which out of the three possible scenarios is true may give a preliminary basis to further hypothesize which evolutionary processes are more likely to dominate the relationship between disease and adaptation genome-wide.

Here, we compare recent adaptation in mendelian disease and non-disease genes in order to disentangle the connections between adaptation and disease. We specifically compare the abundance of recent selective sweeps signals, where hitchhiking has raised haplotypes that carry an advantageous variant to higher frequencies (Smith and Haigh, 1974). Note that this means that we can only compare adaptation at specific loci between disease and non-disease genes that was strong enough to induce hitchhiking, hence we do not take into account polygenic adaptation distributed across a large number of loci that did not leave any hitchhiking signals (see Discussion). As mentioned above, confounding factors may affect the comparison between disease and non-disease genes. In contrast with previous studies, we systematically control for a large number of confounding factors when comparing recent adaptation in human disease and non-disease genes, including evolutionary constraint, mutation rate, recombination rate, the proportion of immune or virus-interacting genes, etc. (please refer to Methods for a full list of the confounding factors included). In addition to controlling for a large number of confounding factors, we estimate false positive risks (FPR) for our comparison pipeline that fully take into account the implications of controlling for many factors (see Methods and Results).

As a list of disease genes to test, we curate human mendelian non-infectious disease genes based on annotations in the DisgeNet and OMIM databases (Methods). We focus on mendelian disease genes rather than all disease genes including complex disease associations, because different evolutionary patterns can be expected between mendelian and complex disease genes based on previous studies (Blekhman et al., 2008; Quintana-murci, 2016; Spataro et al., 2017). In total, we compare 4,215 mendelian disease genes with non-disease genes in the human genome. In agreement with scenario 3, we find a strong deficit of selective sweeps at disease genes compared to non-disease genes. We further test multiple potential explanations for this deficit, and find that higher pleiotropy at disease genes is unlikely to explain the less frequent occurrence of sweeps. In contrast, we find that the sweep deficit at disease genes strongly depends on recombination and the number of known disease variants at given disease genes. This suggests that segregating deleterious mutations at disease genes might interfere with, and slow down genetically linked adaptive variants enough to produce the observed lack of sweeps at disease genes.

## Results

### Controlling for confounding factors with a bootstrap test

To compare disease and non-disease genes, we first ask which potential confounding factors differ between the two groups of genes. As expected, multiple measures of selective constraint are significantly higher in disease compared to non-disease genes. As a measure of long-term constraint, the density of conserved elements across mammals is slightly higher at disease genes compared to non-disease genes (Figure 1: conserved 50kb, conserved 500kb; Methods). As a measure of more recent constraint, we contrast pS, the average proportion of variable synonymous sites, with pN, the average proportion of variable nonsynonymous sites (Figure 1; Methods). If the coding sequences of disease genes are more constrained, we expect a drop of pN at disease genes, but no such drop of pS at neutral synonymous sites. Accordingly, pN is lower at disease compared to non-disease genes, while pS is very similar between the two categories of genes (Figure 1). Therefore, selective constraint was stronger in the coding sequences of disease genes during recent human evolution.

As another measure of recent constraint, we also use McVicker’s B estimator of background selection (McVicker et al., 2009). The amount of background selection at a locus can be used as a proxy for recent constraint, since it depends on the number of deleterious mutations that were recently removed at this locus. The lower B, the more background selection there is at a specific locus. In line with higher recent constraint at disease genes, B is slightly, but significantly lower at disease genes (Figure 1; Methods). Overall, we find evidence of higher constraint at disease genes.

In addition to constraint, mutation rate could represent an important confounder. The proportion of variable neutral synonymous sites pS can be used to compare mutation rates, since the number of variable sites is proportional to the mutation rate under neutrality. As mentioned already, pS is very similar at disease and non-disease genes (Figure 1), suggesting that mutation rates are similar at disease and non-disease genes. This is further supported by the fact that multiple factors that could affect the mutation rate such as GC content or recombination are also similar at disease and non-disease genes (Figure 1; Methods). Aside from mutation rate and constraint, multiple other factors that could affect adaptation differ between disease and non-disease genes, notably including the proportion of genes that interact with viruses, the proportion of immune genes, or the number of protein-protein interactions (PPIs) in the human PPIs network. All these factors have been shown to affect adaptation (Methods), further showing the necessity to control for confounding factors when comparing adaptation at disease and non-disease genes.

### Less sweeps at disease genes

For our comparison of disease and non-disease genes, we measure recent adaptation around human protein coding genes (Methods) using the integrated haplotype score (iHS, (Voight et al., 2006)) and the number of Segregating sites by Length (*nS_L_*, (Ferrer-Admetlla et al., 2014)) in 26 populations (The 1000 Genomes Project Consortium, 2015) (Methods). The iHS and *nS_L_* statistics are both sensitive to recent incomplete sweeps, and have the advantage over other sweep statistics of being insensitive to the confounding effect of background selection (Enard et al., 2014; Schrider, 2020). To evaluate the prevalence of sweeps at disease genes relative to non-disease genes, we do not use the classic outlier approach, and instead used a previously described, more versatile approach based on block-randomized genomes to estimate unbiased false positive risks for whole enrichment curves (Figure 2) (Enard and Petrov, 2020). We first rank genes based on the average iHS or *nS_L_* in genomic windows centered on genes (Methods), from the top-ranking genes with the strongest sweep signals to the genes with the weakest signals. We then slide a rank threshold from a high rank value to a low rank value (from top 5,000 to top 10, x-axis on Figure 2). For each rank threshold, we estimate the sweep enrichment (or deficit) at disease relative to non-disease genes (Figure 2, y-axis). For example, for rank threshold 200, the relative enrichment (or deficit) is the number of disease genes in the top 200 ranking genes, divided by the number of control non-disease genes in the top 200. By sliding the rank threshold, we estimate a whole enrichment curve that is not only sensitive to the strongest sweeps but also to weaker sweeps signals (for example using the top 5,000 threshold; Figure 2). Using block-randomized genomes (Methods), we can then estimate an unbiased false positive risk (FPR) for the whole enrichment curve. This strategy makes less assumptions on the expected strength of selective sweeps. The approach also makes it possible to estimate a single false positive risk based on the cumulated enrichment (or deficit) over multiple whole enrichment curves (Methods). Here, we estimate a single false positive risk for both iHS and *nS_L_* curves considered together, and also for multiple window sizes to measure average iHS and *nS_L_* (from 50kb to 1Mb, Methods).

**Figure 2.**
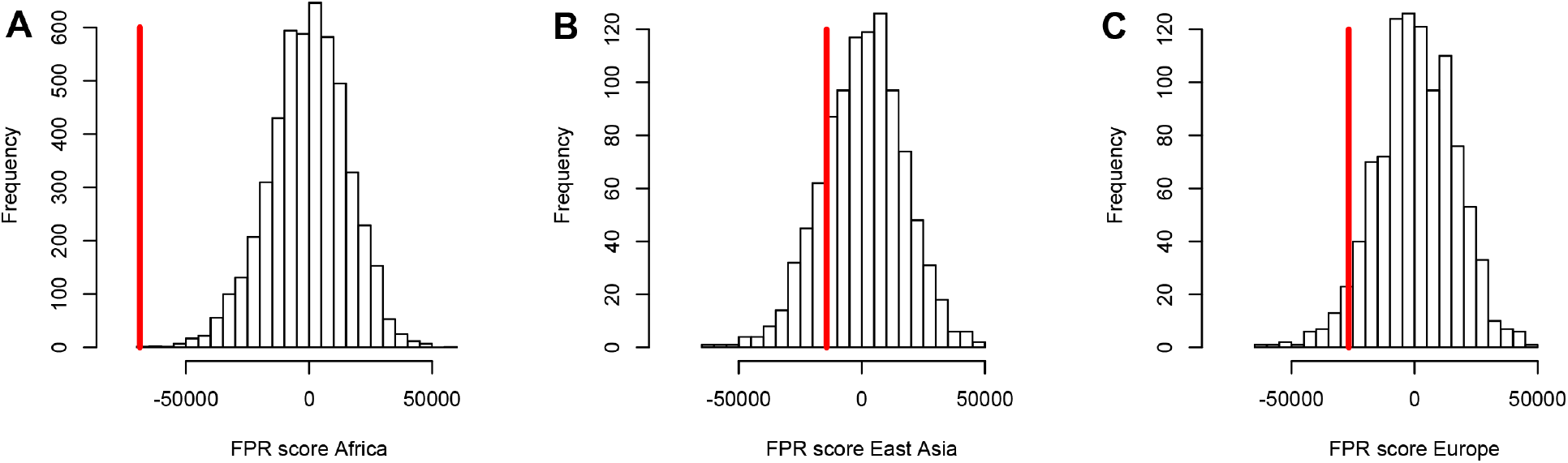
A stronger sweep deficit at disease genes in Africa than in East Asia and Europe. The figure shows the observed sweep enrichment/deficit score used to measure the false positive risk (FPR) in the real genome (red line), compared to the expected null distribution of the score estimated with block-randomized genomes (5,000 block-randomized genomes in Africa, 1,000 in East Asia and Europe; Methods). The FPR score is based on summing the difference between the number of genes in sweeps at disease genes and the number of genes in sweeps in control genes, over both iHS and *nS_L_*, and different window sizes (Methods). A) FPR score in Africa, estimated summing over the ESN, GWD, LWK, MSL and YRI populations from the 1,000 Genomes Project. B) FPR score in East Asia, estimated summing over the CDX, CHB, CHS, JPT and KHV populations. C) FPR score in Europe, summing over the CEU, FIN, GBR, IBS and TSI populations.

To control for confounding factors (Figure 1), we compare sweep signals at disease genes with control non-disease genes that were chosen by a bootstrap test (Castellano et al., 2019; Enard and Petrov, 2020) because they match disease genes in terms of confounding factor values (Methods). Furthermore, control non-disease genes are chosen far from disease genes (>300kb; Methods). We do this to avoid choosing as controls non-disease genes that are too close to disease genes and thus likely to have the same sweep profile (especially in the case of large sweeps potentially overlapping both neighboring disease and non-disease genes). This, together with the large number of confounding factors that we match, tends to limit the pool of possible control genes (Methods). The statistical impact of a limited control pool is however fully taken into account by the estimation of a FPR with block-randomized genomes (Methods).

Because they have experienced different demographic histories, we test different human populations from distinct continents separately. Specifically, we test African populations, East Asian populations and European populations from the 1,000 Genomes Project phase 3 (The 1000 Genomes Project Consortium, 2015). At this stage we must consider the fact that most gene-disease associations in our dataset were likely discovered in European cohorts. Because disease genes in Europe may not always be disease genes in other populations, we cannot exclude the possibility that a sweep enrichment or a sweep deficit might be more pronounced in Europe, unless the evolutionary processes that make a gene more likely to be a disease gene predated the split of different human populations. Conversely, one might expect distinct selective patterns between disease and non-disease genes to be more visible in Africa. Indeed, more intense drift, due to the more severe bottlenecks experienced by ancestral Eurasian populations (The 1000 Genomes Project Consortium, 2015), is expected to dilute true selective patterns among false positive signals more in Europe and East Asia, by creating a higher base level of drift noise.

Using both iHS and *nS_L_* sweep signals, we find a strong depletion in sweep signals at disease genes, especially in Africa with a low false positive risk (FPR=3.10-4 vs. 0.18 in East Asia and 0.05 in Europe, Figure 2A, B and C respectively; Methods). Note that this FPR takes the clustering of multiple genes in the same sweeps into account (Enard and Petrov, 2020). A stronger depletion in Africa suggests that the evolutionary processes linking disease and adaptation at the gene level predate the split of African and European populations, given that most gene-disease associations studies involved European cohorts. The stronger depletion in Africa also suggests that the same pattern might be present outside of Africa, but more hidden by genetic drift noise. It might indeed be harder to distinguish a deficit of true sweep signals at disease genes if it is swamped by an elevated level of false sweep signals occurring at random in the genome, due to more intense drift. Figure 3A, B and C show the sweep deficit curves at disease genes compared to control non-disease genes in Africa, East Asia and Europe, respectively.

**Figure 3.**
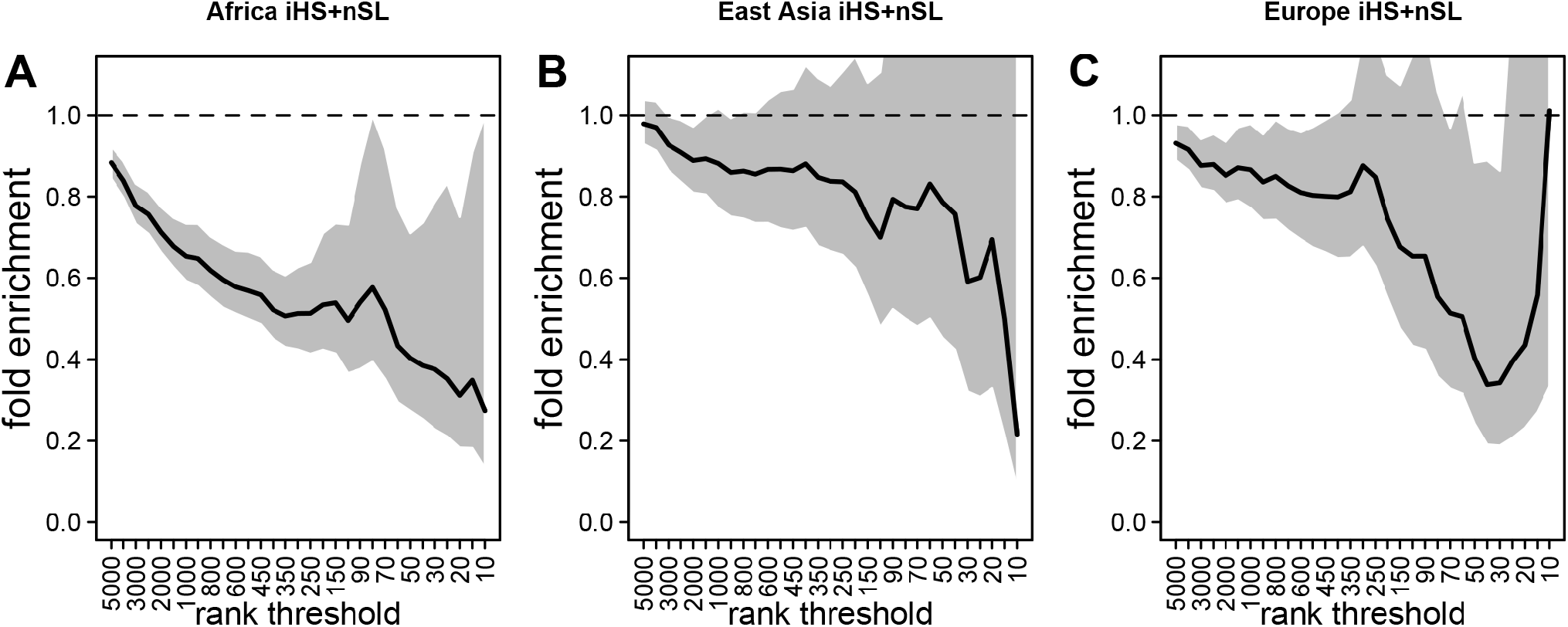
Deficit of iHS and *nS_L_* sweep signals at disease genes. The figure shows the averaged whole enrichment curves and their averaged confidence intervals from the bootstrap test, averaged over both iHS and *nS_L_* sweep ranks, and over all the populations from each continent (Methods). The y-axis represents the relative sweep enrichment at disease genes, calculated as the number of disease genes in putative sweeps, divided by the number of control non-disease genes in putative sweeps. The gray areas are the 95% confidence interval for this ratio. The number of genes in putative sweeps is measured for varying sweep rank thresholds. For example, at the top 100 rank threshold, the relative enrichment is the number of disease genes within the top 100 genes with the strongest sweep signals (either according to iHS or *nS_L_*), divided by the number of control non-disease genes within the top 100 genes with the strongest sweep signals. We use genes ranked by iHS or *nS_L_* using 200kb windows, since 200kb is the intermediate size of all the window sizes we use (50kb, for the smallest, 1000kb for the largest; see Methods). A) Africa, average over the ESN, GWD, LWK, MSL and YRI populations from the 1,000 Genomes Project. B) East Asia, average over the CDX, CHB, CHS, JPT and KHV populations. C) Europe, average over the CEU, FIN, GBR, IBS and TSI populations.

Notably, the stronger depletion observed in Africa likely excludes the possibility that it could be mostly due to a technical artifact, where sweeps themselves might make it harder to identify disease genes in the first place. Sweeps increase linkage disequilibrium (LD) in a way that could make it more difficult to assign a disease to a single gene in regions of the genome with high LD and multiple genes genetically linked to a disease variant. This could result in a depletion of sweeps at monogenic disease genes, simply because disease genes are less well annotated in regions of high LD. However, if this was the case, because most disease gene were identified in Europe, we would expect such an artifact to deplete sweeps at disease genes primarily in Europe, not in Africa. This artifact is also very unlikely due to the fact that recombination rates are similar between disease and non-disease genes (Figure 1). Overall, these results support the third scenario where evolutionary processes decrease adaptation at disease genes. That said, it is important to note that we only detect a deficit of adaptation strong enough to leave hitchhiking signals. Our results do not imply that the same is true for adaptation that is too polygenic to leave signals detectable with iHS or *nS_L_*. Note that the sweep deficit at disease genes in Africa is robust to differences in gene functions between disease and non-disease genes according to a Gene Ontology analysis (Methods) (Gene Ontology Consortium, 2021).

### A limited role of pleiotropy

A deficit of strong adaptation (strong enough to affect iHS or *nS_L_*) raises the question of what creates this deficit at disease genes. Because disease genes tend to be pleiotropic and many disease genes are involved in multiple diseases (see below), pleiotropy is a particularly attractive potential explanation for the lack of sweeps at disease genes. Pleiotropy is defined as the ability for a gene to affect multiple phenotypes. The involvement in multiple phenotypes may make it more difficult for mutations to emerge at pleiotropic genes without any adverse antagonistic effects (Otto, 2004). In addition to the higher selective constraint already accounted for, pleiotropy may thus also make it less likely for advantageous mutations to be advantageous and cause a sweep (Otto, 2004), with the advantage provided by changes at specific phenotypes being mitigated by the adverse effects on other phenotypes.

We can test the involvement of pleiotropy with our dataset by comparing sweeps at disease genes involved in multiple diseases, with sweeps at disease genes involved in only one disease. If pleiotropy decreases the rate of sweeps at disease genes, we predict that genes involved in multiple diseases should experience less sweeps than genes involved in only one disease. There are 1221 disease genes in our dataset associated with five or more diseases (five+ disease genes), and 1296 disease genes associated with only one disease according to the CUI (Concept Unique Identifiers) classification provided by DisGeNet (Methods). When comparing the five+ disease genes with one disease genes far away (>300 kb as when comparing all disease genes with control non-disease genes), we do not find significantly less iHS and *nS_L_* sweep signals at five+ disease genes in Africa (FPR=0.46). This result makes it unlikely that pleiotropy can explain the sweep deficit at disease genes.

### A possible role of interference of deleterious mutations

With pleiotropy likely having a limited role, we further test other possible explanations for the sweep deficit at disease genes. Another possibility is that adaptation may be limited at disease genes due to deleterious mutations interfering with and slowing down advantageous variants. This process has been mostly studied in haploid species (Peck, 1994; Johnson and Barton, 2002; Jain, 2019). In diploid species including humans, recessive deleterious mutations specifically have been shown to have the ability to slow down, or even stop the frequency increase of advantageous mutations that they are linked with (Assaf et al., 2015; Uricchio et al., 2019). Uricchio et al. (2019) in particular found evidence of decreased protein adaptation in the regions of the human genome with strong background selection and low recombination. The majority of disease variants are recessive (Amberger et al., 2019). Thus, if segregating recessive deleterious mutations are more common at disease genes, starting with the known disease variants themselves, then their interference could in theory explain the sweep deficit that we observe. This is true even despite the fact that we matched disease and control non-disease genes for multiple measures of selective constraint. Indeed, we use measures of selective constraint such as the density of conserved elements or the proportion of variable non-synonymous sites pN (Methods), that are indicative of the amount of deleterious mutations that get ultimately removed, but do not provide any detailed information on either the strength of negative selection, or on dominance coefficients. Disease genes and control non-disease genes may have very similar densities of conserved elements and similar pN, and still very different distributions of selection and dominance coefficients of deleterious mutations. Unfortunately, disentangling selection from dominance coefficients is notoriously difficult, because different combinations of selection and dominance coefficients can result in the same patterns of genetic variation (Huber et al., 2018). Although directly comparing the actual total numbers of recessive deleterious mutations at disease and non-disease genes is therefore not possible, we can still use indirect comparison strategies. First, if an interference of deleterious mutations is involved, then this interference is expected to be stronger in low recombination regions of the genome, where more deleterious mutations are likely to be genetically linked to an advantageous mutation. Therefore, we predict that the sweep deficit should be more pronounced when comparing disease and non-disease genes only in low recombination regions of the genome, where the linkage between deleterious and advantageous variants is higher. Conversely, the sweep deficit should be less pronounced in high recombination regions of the genome. Second, if the number of known disease variants at a given disease gene correlates well enough with the total number of segregating recessive deleterious mutations at this disease gene, then we should observe a stronger sweep deficit at disease genes with many known disease variants, compared to disease genes with few known disease variants. Based on these two predictions, the sweep deficit should be particularly strong at disease genes with both many disease variants AND lower recombination. As the number of disease variants for each disease gene, we use the number of disease variants as curated by OMIM/UNIPROT (Methods).

For these comparisons we focus solely on African populations for which we found the strongest sweep deficit (Figure 2). We first compare disease and control non-disease genes both from only regions of the genome with recombination rates lower than the median recombination rate (1.137 cM/Mb). In agreement with recombination being involved, we find that the sweep deficit at low recombination disease genes is much more pronounced than the overall sweep deficit found when considering all disease and control non-disease genes regardless of recombination (Figure 4, FPR=2.10^−4^). Conversely, the sweep deficit at disease genes compared to non-disease genes is much less pronounced when restricting the comparison to genes with recombination rates higher than the median recombination rate (1.137 cM/Mb), and remains only marginally significant (Figure 4, FPR=0.029). This provides evidence that genetic linkage may indeed be involved. Low recombination is however not sufficient on its own to create a sweep deficit, and we further test if the sweep deficit also depends on the number of disease variants at each disease gene. In our dataset, approximately half of all the disease genes have five or more disease variants, and the other half have four or less disease variants (Methods). In further agreement with possible interference of recessive deleterious variants, the sweep deficit is much more pronounced at disease genes with five or more disease variants (Figure 4, FPR=8.10^−4^). The sweep deficit at disease genes with four or less disease variants is barely significant compared to control non-disease genes (Figure 4, FPR=0.032). In addition, disease genes with five or more disease variants, but with recombination higher than the median recombination rate, do not have a strong sweep deficit either (Figure 4, FPR=0.026). A higher number of disease variants alone is thus not enough to explain the sweep deficit. In a similar vein, disease genes with a recombination rate less than the median recombination rate, and with four or less disease variants, do not exhibit a strong sweep deficit (Figure 4, FPR=0.021). This confirms that low recombination alone is not enough to explain the sweep deficit at disease genes. Accordingly, disease genes with both low recombination AND five or more disease variants show the strongest sweep deficit (Figure 4, FPR=2.10^−4^). Disease genes with both high recombination AND less than 5 disease variants show no sweep deficit at all, with a sweep prevalence undistinguishable from control non-disease genes (Figure 4, FPR=0.74). The latter result is important, because it suggests that interference of recessive deleterious variants may be sufficient on its own to explain the whole sweep deficit at disease genes. Both higher linkage and more disease variants seem to be needed to explain the sweep deficit at disease genes. Note that these results are not due to introducing a bias in the overall number of variants by using the number of disease variants, because we always match the level of neutral genetic variation between disease genes and control non-disease genes with pS. The overall level of genetic variation is further matched thanks to pN and thanks to McVicker’s B, whose value is directly dependent on the level of genetic variation at a given locus (McVicker et al., 2009).

**Figure 4.**
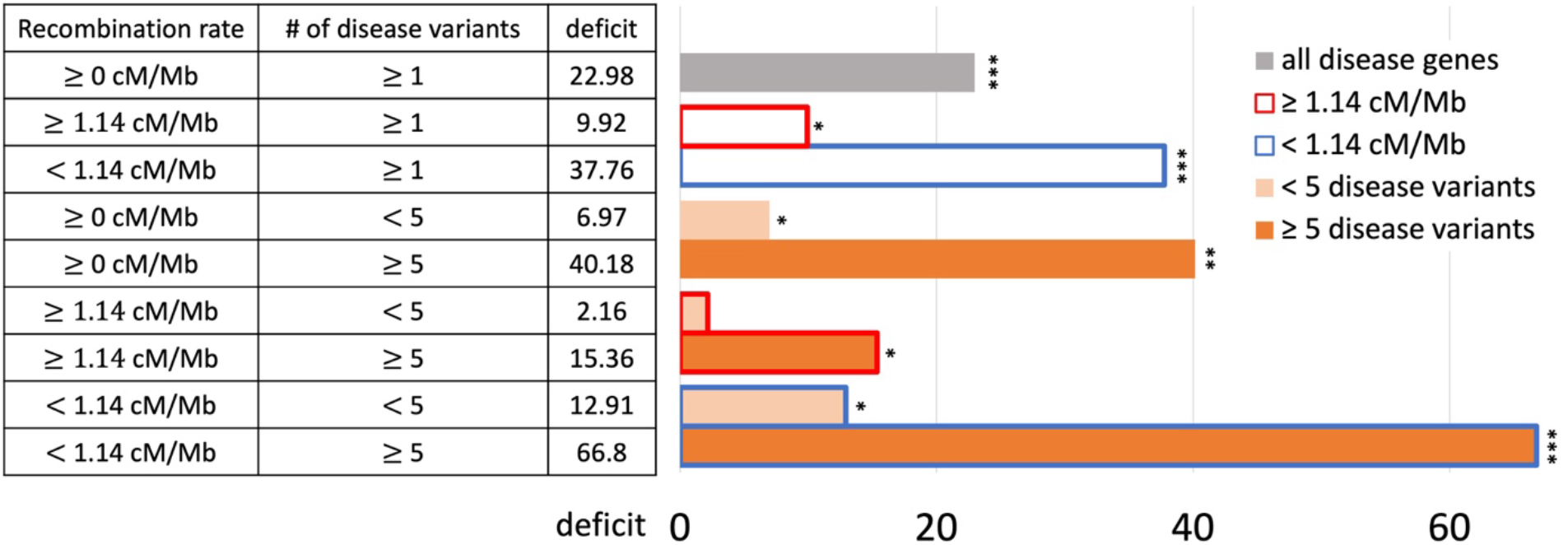
Sweep deficit as a function of recombination and disease variants number. The sweep deficit is measured as the FPR score per gene (to make all tested groups comparable) over all window sizes, and *nS_L_* and iHS, as in Figure 1 (Methods). The different groups are separated according to recombination and numbers of disease variants so that they have approximately the same size (a half or a fourth of the disease genes). All deficits are measured using only African populations.

### Similar levels of sweep depletion in disease genes across MeSH disease classes

Because we found an overall sweep depletion at disease genes, we further ask if genes associated with different diseases might show different patterns of depletion (always in African populations). We classify disease genes into different classes according to the Medical Subject Headings (MeSH) annotation for diseases in DisGeNet (Piñero et al., 2020). The MeSH annotations organize the disease genes into 24 broad disease categories that overlap with distinct organs or large physiological systems (for example the endocrine system). We find significant (FPR<0.05) sweep depletions for all but one disease MeSH classes (FPR<0.05; Figure 5). The sweep deficit is mostly comparable across MeSH disease classes (Figure 5), suggesting that the evolutionary process at the origin of the sweep deficit is not disease-specific. This is compatible with a non-disease specific explanation such as recessive deleterious variants interfering with adaptive variants. The only non-significant deficit is for the MeSH term immune system diseases. Interestingly, there is evidence that past adaptation at disease genes in response to diverse pathogens has resulted in increased prevalence of specific auto-immune diseases (Barreiro and Quintana-Murci, 2010), and we can speculate that this is why we do not see a sweep deficit at those genes.

**Figure 5.**
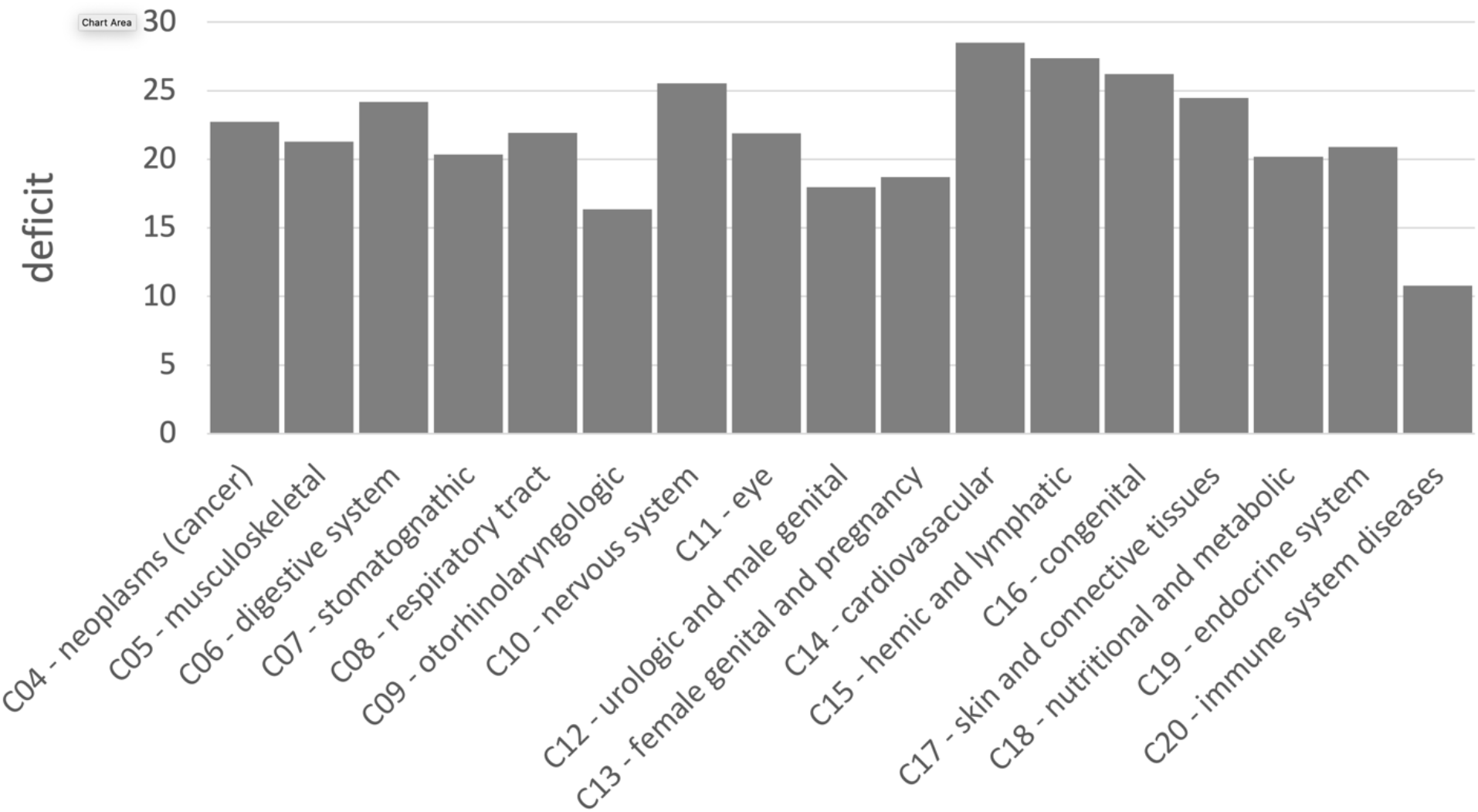
Sweep deficit per MeSH disease classes. The sweep deficit is measured as the overall FPR score per gene (Methods), to make all MeSH classes comparable even if they include different numbers of genes.

## Discussion

We found a depletion of the number of genes in recent sweeps at human non-infectious, mendelian disease genes compared to non-disease genes. Although more work is now needed, the lack of sweeps at disease genes already favors specific evolutionary processes over others. For example, it makes it unlikely that past adaptations increasing the occurrence of disease variants through hitchhiking would be the dominant process linking disease and adaptation at the gene level. The lack of sweeps at disease genes also seems to be unrelated to any difference in mutation accumulation between disease and non-disease genes, since we find no sign of a difference in mutation rates between the two categories of genes in the first place, and since we match metrics accounting for mutation rate in our comparisons (for example, GC content and pS). Instead, a lack of sweeps, once selective constraint has been controlled for, seems to favor a relationship involving a lag of adaptation at disease genes beyond simple constraint (measured by the amount of deleterious mutations that are removed).

Multiple mechanisms might explain such a lag of adaptation. A first possible hypothesis is that disease genes are genes that can be sensitive to the environment and whose fitness optimum can change during evolution when the environment changes. However, when this happens, adaptation then might take more time to chase the new optimum. Although higher pleiotropy is a tempting hypothesis to explain such a lag (Otto, 2004), genes involved in multiple diseases do not have a particularly pronounced sweep depletion compared to genes associated with only one disease. Completely excluding pleiotropy may however require more effort, notably by considering measures of pleiotropy other than the number of diseases a gene has been associated with.

Another hypothesis is that disease genes may have a distribution of deleterious fitness effects that is different from other genes, but that the metrics of constraint that we used do not capture this difference. Specifically, we can imagine a case where disease genes have more currently segregating recessive deleterious variants than other genes, and where selective sweeps are impeded due to the interference of genetically linked recessive deleterious variants. The deleterious effects of these variants can reveal themselves when they hitchhike together with an advantageous variant that is just starting to increase in frequency (Assaf et al., 2015). Accordingly, we find a marked sweep depletion when restricting the comparison to disease and non-disease genes in low recombination regions of the genome and with higher numbers of disease variants (Figure 4). All these comparisons are however indirect, and we do not quantify directly the amount of recessive deleterious mutations at disease or non-disease genes. Further verifying that recessive deleterious mutations impede sweeps more at disease than non-disease genes will require showing that recessive deleterious mutations are indeed more abundant at disease genes, ideally by also estimating dominance coefficients. That said, the majority of disease variants are known to be recessive and using the number of disease variants, as done in the present study, should be a good proxy of the actual number of segregating recessive deleterious mutations. Estimating dominance may prove challenging, since it is difficult to distinguish selection coefficient changes from dominance coefficient changes (Huber et al., 2018). Again, our results provide preliminary evidence to further test in the future.

In addition to suggesting possible explanatory evolutionary scenarios, our results highlight a number of potential limitations and biases that also need to be explored in more detail. First, the lack of sweeps at disease genes suggests the possibility of a technical bias against the annotation of disease genes in sweep regions with high LD, as described in the Results. This bias is unlikely to be the dominant explanation for our results, because then we would expect a stronger sweep deficit at disease genes in Europe than in Africa, given that most disease genes were annotated in Europe. The recombination rate at disease genes is also not different from the recombination rate at non-disease genes (Figure 1). The increase of the sweep deficit when comparing disease and non-disease genes only in low recombination regions (Figure 4), where disease annotation would then be more difficult regardless of overlapping a sweep or not, also suggests that this bias is unlikely. That said, it will still be useful to further investigate in the future how much this potential bias might have contributed to our observations.

Second, even though more intense genetic drift seems a reasonable explanation for the less pronounced sweep deficit at disease genes in Europe and East Asia than in Africa, this claim needs to be further tested, for example with population simulations reproducing past population demographic fluctuations. Such simulations would make it possible to test whether or not past bottlenecks in ancestral Eurasian populations were strong enough to erase the sweep deficit signal at disease genes in East Asia and Europe, by swamping it with random false positive sweep signals.

Further work is also required regarding the connection between the sweep deficit and polygenic adaptation not leaving hitchhiking signals. Our results could be explained by a general lack of adaptation at disease genes, or instead by a different balance between sweeps and polygenic adaptation at disease genes, with less sweeps but more polygenic adaptation that would be less affected by interference with deleterious variants. It may be possible to use recent polygenic adaptation quantification tools such as PALM (Stern et al., 2021) to compare its prevalence at disease and non-disease genes.

Finally, there are multiple directions to further analyze the sweep deficit at disease genes that we have not explored in this manuscript. For instance, analyzing the sweep deficit as a function of the time of onset of diseases (early or late in life), might further provide clues to why the sweep deficit exists in the first place. Preliminary comparison of the sweep deficit at specific MeSH disease classes (Figure 5) with known early (congenital diseases) or mostly late onsets (cancer, cardiovascular) however suggests that the average onset time of diseases might not make much of a difference.

In conclusion, although our analysis reveals a strong deficit of selective sweeps at human disease genes, it also suggests that more work is needed to better understand the evolutionary processes at work, and the biases that may have skewed our interpretations. Despite these limitations, our comparison nevertheless already suggests that specific evolutionary relationships between disease genes and adaptation might be more prevalent than others, especially interference between recessive deleterious and adaptive variants. As an important follow-up question, it may now be important to ask how the sweep deficit at disease genes might have hidden interesting adaptive patterns in previous functional enrichment analyses, especially in gene functions that are often annotated based on disease evidence in the first place. For example, metabolic genes are believed to be of particular interest for adaptation to climate change. But metabolic genes are often found due to their role in metabolic disorders, and a strong representation of disease genes among all metabolic genes could then in theory mask any sweep enrichment. A sweep enrichment at metabolic genes might only become visible once controlling for the proportion of disease genes, in addition to the list of controls that we already use in the present analysis (Methods). Our results thus highlight the complexity of studying functional patterns of adaptation in the human genome.

## Methods

### Disease gene lists

We consider genes that are known to be associated with diseases as disease genes. We focus on protein-coding genes associated with human mendelian non-infectious diseases. Complex diseases are associated with several loci and environmental factors. Patterns of positive selection at complex disease and mendelian disease genes may differ (Blekhman et al., 2008), which is why we restrict our analysis to mendelian disease genes. We also restrict our analyses to non-infectious disease genes, since interactions with pathogens are an entirely different problem. We nevertheless control for the proportion of genes that are immune genes or interact with viruses (see below), since it has been shown that immune genes and interactions with viruses drive a large proportion of genomic adaptation in humans (Enard et al., 2016; Castellano et al., 2019). Therefore, different proportions of immune and virus-interacting genes between disease and non-disease genes might confound their comparison. Moreover, although diseases can be associated with non-coding genes, we only use protein-coding genes. We curate disease genes defined as genes associated with diseases according to both DisGeNet (Piñero et al., 2020) and OMIM (Amberger et al., 2019), to ensure that we focus on high-confidence disease genes. DisGeNet is a comprehensive database including gene-disease associations (GDAs) from many sources. In order to get disease genes with high confidence, we further only use GDAs curated by UniProt. These gene-disease associations are extracted and carefully curated from the scientific literature and the OMIM (Online Mendelian Inheritance in Man) database, which reports phenotypes either mendelian or possibly mendelian (Amberger et al., 2019). We also exclude all genes associated with infectious diseases according to MeSH annotation (disease class C01). In the end, we curate 4215 non-infectious mendelian disease genes from DisGeNet also curated by OMIM and Uniprot. Although we rely on GDAs from Uniprot to curate high-quality disease genes, we also include GDAs of DisGeNet from other sources when classifying disease genes into different MeSH classes and measuring pleiotropy, as long as a disease gene has at least one GDA curated by OMIM and Uniprot. We completely exclude GDAs that are only reported by CTD (Comparative Toxicogenomics Database) (Davis et al., 2021) in this study. This is because CTD includes a broad range of chemical-induced diseases that might only happen where people are exposed to these chemicals, especially some inorganic chemicals that may not be present in natural environments (Davis et al., 2021).

In order to study different types of diseases, we also divide disease genes into different classes according to the annotated MeSH classes in DisGeNet (Piñero et al., 2020). Those diseases without MeSH class are annotated as “unclassfied”. Genes belonging to more than one MeSH class are counted in each MeSH class where they are present. MeSH classes including less than 50 genes are not considered in this study. We classify all the non-infectious disease genes into 22 MeSH classes including Neoplasms (C04), Musculoskeletal Diseases (C05), Digestive System Diseases (C06), Stomatognathic Diseases (C07), Respiratory Tract Diseases (C08), Otorhinolaryngologic Diseases (C09), Nervous System Diseases (C10), Eye Diseases (C11), Male Urogenital Disease (C12), Female Urogenital Diseases and Pregnancy Complications (C13), Cardiovascular Diseases (C14), Hemic and Lymphatic (C15), Congenital, Hereditary, and Neonatal Diseases and Abnormalities (C16), Skin and Connective Tissue Diseases (C17), Nutritional and Metabolic Diseases (C18), Endocrine System Diseases (C19), Immune System Diseases (C20), Mental Disorders (F03) and “unclassified”.

### Detecting selection signals at human genes

All the analyses were conducted human genome version hg19. We use two different methods to detect selective sweeps in human populations: iHS (integrated Haplotype Score, Voight et al., 2006) and *nS_L_* (Ferrer-Admetlla et al., 2014). Both approaches are haplotype-based statistics calculated with polymorphism data. We use human genome data from the 1,000 Genomes Project phase 3, which includes 2,504 individuals from 26 populations (The 1000 Genomes Project Consortium, 2015).

We measure iHS and *nS_L_* in windows centered on human coding genes (i.e. windows whose center is located half-way between the most upstream transcript start site and most downstream transcript stop site of protein coding genes). We use windows of sizes ranging from 50 kb to 1,000 kb (50kb, 100kb, 200kb, 500kb and 1,000kb) since we do not want to presuppose of the size of sweeps, and since the size of the selective sweeps may vary between different genes. Moreover, to avoid any preconception related to the expected strength or number of sweep signals, we use a moving rank threshold strategy to measure the enrichment or deficit in sweeps at disease genes. For example, we select the top 500 genes with the stronger sweep signals according to a specific statistic (iHS or *nS_L_*). We then compare the number of diseases and non-disease genes within the top 500 genes with the strongest iHS or *nS_L_* signals. This was repeated for different top thresholds and the corresponding ranks from top 5,000 to top 10 (Figure 3). Genes are ranked based on the average iHS or *nS_L_* in their gene centered windows. Both iHS and *nS_L_* measure, individually for each SNP in the genome, how much larger haplotypes linked to the derived SNP allele are compared to haplotypes linked to the ancestral allele (Voight et al., 2006; Ferrer-Admetlla et al., 2014). For each window, we measure the average of the absolute value of iHS or *nS_L_* over all the SNPs in that window with an iHS or *nS_L_* value. The average iHS or *nS_L_* values in a window provide high power to detect recent select sweeps (Enard and Petrov, 2020).

### Comparing recent adaptation between disease and non-disease genes

We use a previously developed gene-set enrichment analysis pipeline to compare recent adaptation between disease and non-disease genes (Enard and Petrov, 2020) (https://github.com/DavidPierreEnard/Gene_Set_Enrichment_Pipeline). This pipeline includes two parts. The first part is a bootstrap test that estimates the whole sweep enrichment or depletion curve at genes of interest (disease genes in our case). The second part is a false positive risk (also known as false discovery rate in the context of multiple testing) that estimates the statistical significance of the whole sweep enrichment curve using block-randomized genomes.

To compare disease and non-disease genes, we first need to select control non-disease genes that are sufficiently far away from disease genes. In that way, we avoid using as controls non-disease genes that overlap the same sweeps as neighboring disease genes, thus resulting in an underpowered comparison. The question is then how far do we need to choose non-disease control genes? Ideally, we would choose non-disease control genes as far as possible from disease genes in the human genome, further than the size of the largest known sweeps (for example the lactase sweep), which would be on the order of a megabase. However, because there are many disease genes in our dataset (4,215), there are very few non-disease genes in the human genome that are more than one megabase away from the closest disease gene. This is a problem, because the available number of potential control non-disease genes is an important parameter that can affect both the type I error, false positive rate, and type II error, false negative rate of the disease vs. non-disease genes comparison. Indeed, the smaller the control set, the more likely it is to deviate from being representative of the true null expectation at non-disease genes. The noise associated with a small sample could go either way. Either the small control sample happens by chance to have less sweeps, and the bootstrap test we use to compare disease and non-disease genes will become too liberal to detect sweep enrichments, and to conservative to detect sweep deficits. Or the small control sample happens by chance to have more sweeps than a larger control sample would, and the bootstrap test becomes too conservative to detect sweep enrichments, and too liberal to detect sweep deficits.

After trying distances between disease genes and control disease genes of 100kb, 200kb, 300kb, 400kb and 500kb, we find that the sweep deficit observed at disease genes increases steadily from 100kb to 300kb (Table 1), showing that 100kb or 200kb are likely insufficient distances. Further than 300kb at 400kb, we do not observe much stronger sweep deficits than at 300kb, while at the same time the risks of type I and type II errors keep increasing due to shrinking non-disease genes control sets. This would translate in a decreased power to possibly exclude the null hypothesis of no sweep enrichment or deficit in the second part of the pipeline, when estimating the actual pipeline FPR. Because of this, we set the required distance of potential control non-disease genes from disease genes at 300kb. This is also the distance where there are still approximately as many control genes (3455) as there are disease genes that we can use for the comparison (3030; those genes out of the 4,215 disease genes with sweep data and data for all the confounding factors).

**Table 1.** Sweep deficit as a function of the minimal distance of control non-disease genes. The sweep deficit is measured by the FPR score, that is the cumulative difference between the number of genes in sweeps at disease and control non-disease genes, across window sizes, sweep summary statistics, and African populations (see the rest of the Methods).

| minimal distance | sweep deficit |
| --- | --- |
| 100kb | -20889 |
| 200kb | -35009 |
| 300kb | -68928 |
| 400kb | -88546 |

Another important aspect of the bootstrap test (first part of the pipeline), aside from setting up the minimal distance of the control non-disease genes, is the matching of potential confounding factors likely to influence sweep occurrence. We choose non-disease control genes that have the same confounding factors characteristics as disease genes (for example, control non-disease genes that have the same gene expression level across tissues as disease genes). The precise matching algorithm is detailed in Enard & Petrov (2020).

When comparing disease and non-disease genes with the bootstrap test, we control for the following potential confounding factors that could influence the occurrence of sweeps at genes:

- Average overall expression in 53 GTEx v7 tissues (The GTEx Consortium, 2015) (https://www.gtexportal.org/home/). We used the log (in base 2) of TPM (Transcripts Per Million).
- Expression (log base 2 of TPM) in GTEx lymphocytes. Expression in immune tissues may impact the rate of sweeps.
- Expression (log base 2 of TPM) in GTEx testis. Expression in testis might also impact the rate of sweeps.
- deCode recombination rates 50kb and 500kb: recombination is expected to have a strong impact on iHS and *nS_L_* values, with larger, easier to detect sweeps in low recombination regions but also more false positive sweeps signals. The average recombination rates in the gene-centered windows are calculated using the most recent deCode recombination map (Halldorsson et al., 2019). We use both 50kb and 500kb window estimates to account for the effect of varying window sizes on the estimation of this confounding factor (same logic for other factors where we also use both 50kb and 500kb windows).
- GC content is calculated as a percentage per window in 50kb and 500kb windows. It is obtained from the USCS Genome Browser (Kent et al., 2002).
- The density of coding sequences in 50kb and 500kb windows centered on genes. The density is calculated as the proportion of coding bases respect to the whole length of the window. Coding sequences are Ensembl v99 coding sequences.
- The density of mammalian phastCons conserved elements (Siepel et al., 2005) (in 50kb and 500k windows), downloaded from the UCSC Genome Browser (Kent et al., 2002). We used a threshold considering 10% of genome as conserved, as it is unlikely that more than 10% of the whole genome is constrained according to previous evidence (Siepel et al., 2005). Given that each conserved segment had a score, we considered those segments above the 10% threshold as conserved.
- The density of regulatory elements, as measured by the density of DNASE1 hypersensitive sites (in 50kb and 500kb windows) also from the UCSC Genome Browser (Kent et al., 2002).
- The number of protein-protein interactions (PPIs) in the human protein interaction network (Luisi et al., 2015). The number of PPIs has been shown to influence the rate of sweeps (Luisi et al., 2015). We use the log (base 2) of the number of PPIs.
- The gene genomic length, i.e. the distance between the most upstream and the most downstream transcription start sites.
- The number of gene neighbors in a 50kb window, and the same number in 500kb window centered on the focal genes: it is the number of coding genes within 25kb or within 250kb.
- The number of viruses that interact with a specific gene (Enard and Petrov, 2020).
- The proportion of immune genes. The matched control sets have the same proportion of immune genes as disease genes, immune genes being genes annotated with the Gene Ontology terms GO:0002376 (immune system process), GO:0006952 (defense response) and/or GO:0006955 (immune response) as of May 2020 (Gene Ontology Consortium, 2021).
- The average number of non-synonymous variants PN in African populations, and the number of synonymous variants PS. We matched PN to build control sets of non-disease genes with the same average amount of strong purifying selection as disease genes. Also, PS can be a proxy for mutation rate and we can build control sets of non-disease genes with similar level of mutation rates.
- McVicker’s B value which can be used to account for the effect of background selection on rates of adaptation and especially weak adaptation (McVicker et al., 2009).

Similar to the selection of control genes far enough from disease genes, the matching of many confounding factors decreases the number of non-disease genes that can effectively be used as controls. This further increases the risk of type I and type II errors of the bootstrap test, as previously described. In addition, the bootstrap test only provides p-value for each tested sweep rank threshold separately, in the whole enrichment (or deficit) curve (Figure 2). It does not provide any estimate of the significance of the whole curve, which is needed to estimate the significance of a sweep enrichment or deficit without making too many assumptions on how many sweeps are expected or how strong they are.

To address the increased type I and type II error risks of the bootstrap test, as well to get an unbiased significance estimate for whole enrichment curves, the second part of our pipeline conducts a false positive risk analysis based on block-randomized genomes (Enard and Petrov, 2020). Briefly, we re-estimate many whole enrichment curves reusing the same disease and control non-disease genes used in the first part of the pipeline by the bootstrap test, but after having randomly shuffled the locations of genes or clusters of neighboring genes in sweeps at those disease and control non-disease genes. To do this, we order the disease and control non-disease genes as they appear in the genome. We then define blocks of neighboring genes, whose limits do not interrupt clusters of genes in the same putative sweep. Then, we randomly shuffle the order of these blocks. Because we do not cut any cluster of genes that might be in the same sweep, the resulting block-randomized genomes preserve the same clustering of the genes in the same putative sweeps as in the real genome. With this approach, we look at the exact same set of disease and control non-disease genes and just shuffle sweep locations between them. Thus, by using many block-randomized genomes, we can estimate the null expected range of whole enrichment curves while fully accounting for the extra variance expected from having a limited sample of control non-disease genes. We can then estimate a false positive risk (FPR) for the whole enrichment or deficit curve by comparing the real observed one with the distribution of random curves generated with block-randomized genomes.

To measure the FPR for a curve, we need to define a metric to compare the real curve with the randomly generated ones. In figure 1, we show relative enrichments at each sweep rank threshold, the number of disease genes in sweeps divided by the number of control non-disease genes in sweeps. As a summary metric for the curve, we could then use the sum of the relative enrichments over all thresholds. However, the issue with this approach is that a relative enrichment is the same whether we have 2 disease genes in sweeps and one control non-disease gene in sweeps, or we have 200 disease genes in sweeps and 100 control non-disease genes in sweeps. Thus, although relative enrichments are convenient for visualization on a figure, they are not adequate to measure the FPR. Instead of the relative enrichment, we use the difference between disease and non-disease genes, that is, the number of disease genes in sweeps, minus the average number of control non-disease genes across control sets built by the bootstrap test. We then use as a metric for a whole curve the sum of differences over all the rank thresholds. We use this sum of differences to estimate the enrichment or deficit curve FPR, as the proportion of block-randomized genomes where the sum of differences exceeds the observed sum of differences for an enrichment (one minus this proportion for a deficit).

Importantly, although so far we have described the case where we measure the FPR for one enrichment curve, nothing prevents us from calculating a single sum of differences over an entire group of enrichment or deficit curves. This way, we can measure a single FPR for any number of curves considered together. In our analysis, we measure a single FPR adding iHS and *nS_L_* curves together, and also adding together the curves for 50kb, 100kb, 200kb, 500kb and 1000kb windows (ten curves in total, 2 statistics*5 window sizes).

### Sweep deficit at high and low recombination disease genes, and at high and low disease variant number disease genes

To generate Figure 4, we separate disease genes in groups of approximately the same size based on their recombination rate and numbers of disease variants annotated in OMIM/Uniprot. We separate the disease genes into two groups of equal size, those with recombination lower than 1.137 cM/Mb, and those with recombination higher than this value. To count the disease variants at each disease gene, we count not only the OMIM/Uniprot disease variants for that gene, but also all the other OMIM/Uniprot disease variants that occur in a 500kb window centered on that gene. We do this because the recessive deleterious variants form other nearby disease genes may also interfere with adaptation. Half of disease genes have less than five OMIM/Uniprot disease variants, and half have five or more.

### Impact of functional differences between disease and non-disease genes on the sweep deficit

The sweep deficit at disease genes could be due to a different representation of gene functions at disease genes compared to control non-disease genes. In this case, disease genes would have less adaptation not because they are disease genes, but because the gene functions that are enriched among disease genes compared to non-disease happen to experience less adaptation. We can test this possibility using Gene Ontology (GO) (Gene Ontology Consortium, 2021) functional annotations as follows. If GO gene functions that are enriched in disease genes experience less adaptation independently of the disease status of genes, then we can predict that non-disease genes with these functions should also experience less adaptation than non-disease genes that do not have these GO functions. In total, we find that 3,097 GO annotations are enriched in disease genes compared to confounding factors-matched controls (bootstrap test P≤0.01). In our dataset, half of non-disease genes have 20 or more of these GO annotations, and half have less than twenty (very few have none). We find no difference in the sweep prevalence between the two groups (20 or more annotations vs. less than 20 annotations at least 300kb away; FPR=0.15). The sweep deficit at disease genes is therefore unlikely to be due to the gene functions that are more represented in disease genes compared to controls. In addition, such a scenario would not explain the lack of sweep deficit observed at disease genes with high recombination rates and low numbers of disease variants (Figure 4).

## Acknowledgements

We wish to thank Dan Shrider for helpful comments on the results presented in the manuscript.

## Author Contributions

Conceived and designed the analyses: CD, DE. Performed the analyses: CD and DE. Wrote the manuscript: CD, DST and DE. Interpreted the results: CD, DST, MEL and DE.

## Notes

### Competing Interest Statement

The authors have declared no competing interest.

